# Sub-second Dopamine Signals during Risky Decision-Making in Patients with Impulse Control Disorder

**DOI:** 10.1101/2023.09.11.557178

**Authors:** Brittany Liebenow, Thomas Wilson, Benjamin Maas, Emily Aladnani, Rosalyn J. Moran, Jason White, Terry Lohrenz, Ihtsham ul Haq, Mustafa S. Siddiqui, Adrian W. Laxton, Stephen B. Tatter, P. Read Montague, Kenneth T. Kishida

## Abstract

**Background:** Impulse Control Disorder (ICD) in Parkinson’s disease is a behavioral addiction arising secondary to dopaminergic therapies, most often dopamine receptor agonists. Prior research implicates changes in striatal function and heightened dopaminergic activity in the dorsal striatum of patients with ICD. However, this prior work does not possess the temporal resolution required to investigate dopaminergic signaling during real-time progression through various stages of decision-making involving anticipation and feedback.

**Methods:** We recorded high-frequency (10Hz) measurements of extracellular dopamine in the striatum of patients with (N=3) and without (N=3) a history of ICD secondary to dopamine receptor agonist therapy for Parkinson’s disease symptoms. These measurements were made using carbon fiber microelectrodes during awake DBS neurosurgery and while participants performed a sequential decision-making task involving risky investment decisions and real monetary gains and losses. Per clinical standard-of-care, participants withheld all dopaminergic medications prior to the procedure.

**Results:** Patients with ICD invested significantly more money than patients without ICD. On each trial, patients with ICD made smaller adjustments to their investment levels compared to patients without ICD. In patients with ICD, dopamine levels rose or fell on sub-second timescales in anticipation of investment outcomes consistent with increased or decreased confidence in a positive outcome, respectively; dopamine levels in patients without ICD were significantly more stable during this phase. After outcome revelation, dopamine levels in patients with ICD rose significantly more than in inpatients without ICD for better-than-expected gains. For worse-than-expected losses, dopamine levels in patients with ICD remained level whereas dopamine levels in patients without ICD fell.

**Conclusion:** We report significantly increased risky behavior and exacerbated phasic dopamine signaling, on sub-second timescales, anticipating and following the revelation of the outcomes of risky decisions in patients with ICD. Notably, these results were obtained when patients who had demonstrated ICD in the past but were, at the time of surgery, in an off-medication state. Thus, it is unclear whether observed signals reflect an inherent predisposition for ICD that was revealed when dopamine receptor agonists were introduced or whether these observations were caused by the introduction of dopamine receptor agonists and the patients having experienced ICD symptoms in the past. Regardless, future work investigating dopamine’s role in human cognition, behavior, and disease should consider the signals this system generates on sub-second timescales.

## Introduction

Impulse control disorder (ICD) is characterized by a rapid onset of repeated risky decision-making above the norm for an individual that causes significant distress in daily life.^1,2^ ICDs are most often triggered by the start of dopamine receptor agonist therapy, and are commonly observed in the treatment of Parkinson’s disease (PD) where dopaminergic therapies are a routine intervention.^2–4^ Pathologic gambling, excessive shopping, excessive sexual behavior, and binge eating are the most thoroughly documented ICDs.^2,3^ These behaviors comprise a larger class of behavioral addictions (BAs), with ICD in PD being a special case of BAs typically induced by dopaminergic therapies.^2,4,5^ Dopaminergic action triggers the rapid onset of the behavioral addiction symptoms that define ICD.^4,6,7^ The over-activation of dopaminergic systems is also posited to play a role in substance-based, as well as behavior-based, addictions.^6–8^ Studying ICD and its underlying mechanisms may be a model for broader human research on addiction disorders.

Our understanding of dopamine’s role in ICD is founded primarily on non-invasive neuroimaging studies that use indirect methods for assessing the dopaminergic state. A study assessing the risk of ICD with dopamine transporter imaging (DAT-SPECT) data identified lower DAT availability in regions across the striatum as a risk factor for ICD.^9^ Positron emission tomography (PET) imaging results highlight the striatum as a potential region for dopaminergic differences that underlie ICD.^10^ The striatum is a key region in the associative cortico-striatal circuit believed to control behavioral inhibition and impulsivity.^11^ The exact neuroanatomic location of dopaminergic action has not been identified, though there are promising culprit brain regions with contrasting roles in its dysfunction.^11,12^ The caudate and anterior putamen may have heightened dopaminergic activity,^10,11^ whereas the subthalamic nucleus may become silent in patients with ICD; both structures may mediate signals from the orbitofrontal cortex, with a network system failure potentially resulting in increased impulsivity.^11,12^ That dopaminergic action begets the onset of rapid behavioral addiction suggests the possibility that an aberrant dopaminergic system may contribute to impulsivity.^4,6,8,13^

A prior limitation of literature investigating dopamine’s role in impulsivity has been the lack of direct measurements of dopamine signaling on time scales relevant to the speed of human decision-making^14–17^ within patients with ICD. Recently, technology that allows safe measurements of dopamine release with sub-second temporal resolution in humans revealed changes in dopamine levels within hundreds of milliseconds of actions and events around risky decisions and their outcomes.^14–18^ This approach uncovered novel characterization in behavioral and dopaminergic patterns in individuals with alcohol use disorder (AUD),^19^ which is both a comorbidity and clinical predictor of ICD.^3,4,20–22^ In fact, a family history of alcohol disorders is a leading clinical predictor of ICD risk supported by several studies.^3,4,19–21^ One study of 3,090 PD patients found a statistically significant increase in prevalence in family history of alcohol disorders in first-degree relatives in ICD versus non-ICD patients.^3^ A coinciding family or personal history of disordered alcohol or substance use has also been reported in around 28-30% of ICD patients.^3,4^ Further, single photon emission computed tomography (SPECT) neuroimaging evidence demonstrates that ICD patients have lower dopamine transporter (DAT) binding in the striatum consistent with the effects of alcohol,^22^ suggesting a shared underlying dopaminergic mechanism for the symptoms of both addictions. Dopaminergic fluctuations serve as a bridge between behavioral addictions in ICD and the substance-based AUD; investigating underlying dopaminergic fluctuations in humans with and without ICD may allow us to define dopaminergic signatures underlying behavioral patterns for future investigation in humans and other models of addiction. Here, we build on this recent work in AUD^19^ and apply a human voltammetry approach to investigate behavioral and dopaminergic fluctuation patterns in ICD.

The present study investigates differences in dopamine signaling and behavior in patients with ICD using direct measurements from the human brain in the operating room while patients perform a behavioral task (Figure 1A). Dopamine measurements are completed during deep brain stimulation (DBS) electrode implantation surgeries that provide the opportunity to make real-time dopamine measurements while simultaneously collecting decision-making data. The trajectories of the clinical electrodes allow for research recordings in the striatum (i.e., caudate and putamen, Figure 1B and Supplemental Table 1) allowing for measurements directly from hypothesized brain regions of dopaminergic dysfunction in ICD.^9–11^ We compare behavior and associated sub-second dopamine fluctuations in patients with PD with and without a history of ICD secondary to dopamine receptor agonist therapy.

**Figure 1.**
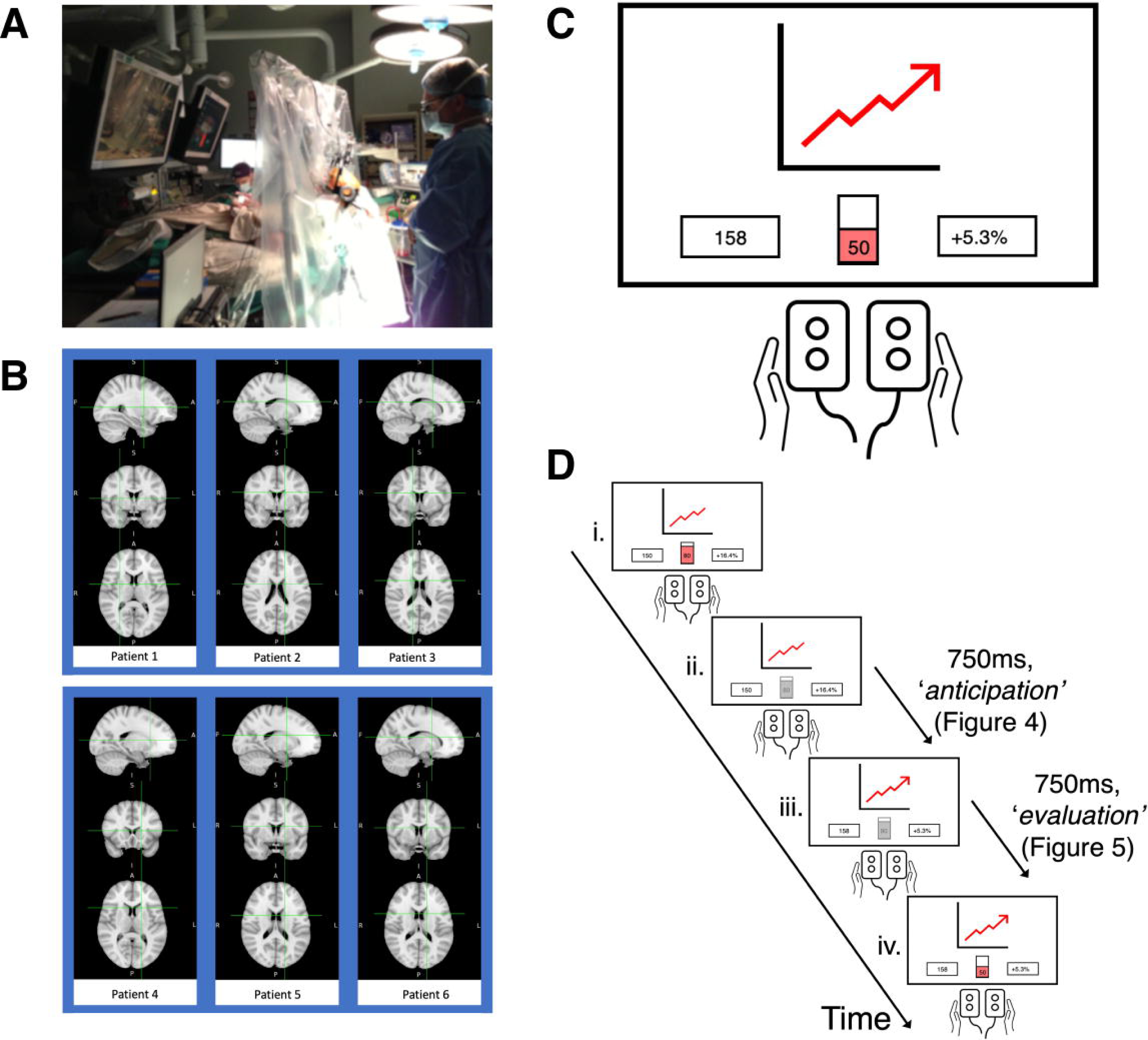
Timeline of events during the ‘stock market investment’ task. Patients undergoing DBS-electrode implantation performed a ‘stock market investment’ task, which is an interactive, sequential, decision-making task. (**A)** Neurosurgical setting: a patient performing the stock market investment task while dopamine measurements are recorded. Participants make decisions using button boxes based on information presented on a computer screen. (**B)** dopamine recording locations for six patients (Patient 1-3: ICD, Patient 4-6: non-ICD). **(C)** The layout of the task on the computer screen is consistent throughout the experiment. The top center box displays a historic stock market chart. The left bottom box shows the participants current portfolio value. The center bottom box (red) shows the participant investment level as a percentage of their current portfolio value; this is a slider bar that can be adjusted in increments of 10%. The bottom right box shows the participant’s most recent return on their investment. (**D)** Timeline of a single trial (120 trials per participant): i. Participants use button boxes to make investment decisions (e.g. 80% of their portfolio value is currently selected). ii. After submitting investment, there is a 750ms delay before the market return is revealed. iii. The outcome of the participant’s investment is displayed simultaneously with the market outcome. A 750ms waiting period occurs before participants can readjust their investment level. iv. Participants use button boxes to make investment decisions (e.g. the participant has decreased their investment to 50% of their portfolio value).

## Materials and methods

### Participant Recruitment

All participant recruitment, consent, and experimental procedures were approved by the institutional review boards of both the Wake Forest University School of Medicine and Virginia Tech. Participant recruitment and informed consent was performed after patients consented to receiving Deep Brain Stimulation (DBS) electrode implantation surgeries for therapeutic intervention. We separate dopamine measurements from PD patients by a history of ICD symptoms (ICD) or no history of ICD symptoms (“non-ICD”, N=3) by physician documentation in the medical record. Patients with ICD (N=3) were identified from a previously collected data set.^16,23^ The non-ICD group (N=3) was selected from the same data set but matched to the ICD group by age and sex (Table 1).

**Table 1.** Clinical characteristics for the ICD and non-ICD groups. This table describes patients (one in each column) by clinical characteristics (one in each row). Patients in the ICD group include patients 1-3. Patients in the non-ICD group include patients 4-6. Answers to binary questions are indicated by a “1” for “yes” and a “0” for “no”. Information related to ICD onset and qualitative reports of symptoms include Impulsivity Type, Impulsivity Resolved with Removal / Reduction of Drug, and Medication Inducing Impulsivity. Surgical information includes Reason for Surgery, Age at Time of Surgery, and Electrochemical Target. Other past medical history includes Antidepressants at Time of Surgery, Dyskinesia, Bradykinesia, Dystonia, Smoking History, and Alcohol Use.

**Clinical Characteristics of Patients with and without Impulse Control Disorder**
| Patient # | 1 | 2 | 3 | 4 | 5 | 6 |
| --- | --- | --- | --- | --- | --- | --- |
| Impulsivity Type | Gambling | Increased purchasing and watching pornography | Increased libido (including unusually rough behavior during intercourse), disinhibited social behavior | None | None | None |
| Impulsivity Resolved with Removal / Reduction of Drug | 1 | 1 | 1 | NA | NA | NA |
| Medication Inducing Impulsivity | Mirapex (Pramipexole) | Mirapex (Pramipexole) | Requip (Ropinirole HCl) | NA | NA | NA |
| Reason for Surgery | PD | PD | PD | PD | PD | PD |
| Age at Time of Surgery | Early 40s | Early 60s | Early 50s | Early 60s | Early 50s | Early 50s |
| Electrochemical Target | Putamen | Caudate | Caudate | Caudate | Caudate | Caudate |
| Antidepressants at Time of Surgery | None | Nortriptyline | None | Escitalopram | Bupropion | None |
| Dyskinesia | 1 | 0 | 1 | 0 | 1 | 0 |
| Bradykinesia | 1 | 1 | 1 | 1 | 1 | 1 |
| Dystonia | 1 | 0 | 0 | 0 | 1 | 0 |
| Smoking History | 0 | 0 | 1 | 1 | 1 | 0 |
| Alcohol Use | 0 | 0 | 0 | 0 | 1 | 0 |

### Task Design (‘Stock Market Investment’ Task)

The ‘stock market investment’ task is a sequential-choice gambling task previously published (Figures 1C and 1D).^14,16,23,24^ Participants view this task on a computer monitor that hangs from the ceiling in the operating room (Figure 1A). Participants start the game with $100 that they are instructed to invest against historical stock market returns that are presented to the participants as stock market charts. Participants are instructed to invest a percentage of their portfolio on each trial (Figures 1C and 1D). Once they submit their investment decision the slider bar is locked out for 750ms (‘anticipation’ phase shown in Figure 1D-ii); then the next market section is revealed along with the participant’s return (based on the market change and their invested amount); the slider bar remains locked out for an additional 750ms (‘evaluation’ phase Figure 1D-iii) before the participant can begin the next trial by readjusting their investment and eventually submitting their next investment decision.^14,16,23,24^

### Behavioral Analysis of Investment Behavior

We calculated the average investment for each group by first averaging the investments for each participant (N=3 for ICD and N=3 for non-ICD). We created group averages by averaging the individual-level average investments based on ICD status (Figure 2A). We sorted investments for each group by first pooling the investments for all participants in a single group. We separated the distribution of investments by ICD group (Figure 2B). We calculated the average change in investment for each group separated by decreasing and increasing investments. Increasing investments were defined as investments with a change from the prior trial of greater than 0%. Decreasing investments were defined as investments with a change from the prior trial of less than 0%. We calculated the average increasing and decreasing investment changes by first averaging in the investment changes for each individual. We created group averages by averaging the individual-level increasing and decreasing investment changes based on ICD status (Figure 3A). We sorted investment changes for each group by first pooling the investment changes for all participants in a single group. We separated the distribution of investment changes by ICD group (Figure 3B).

**Figure 2.**
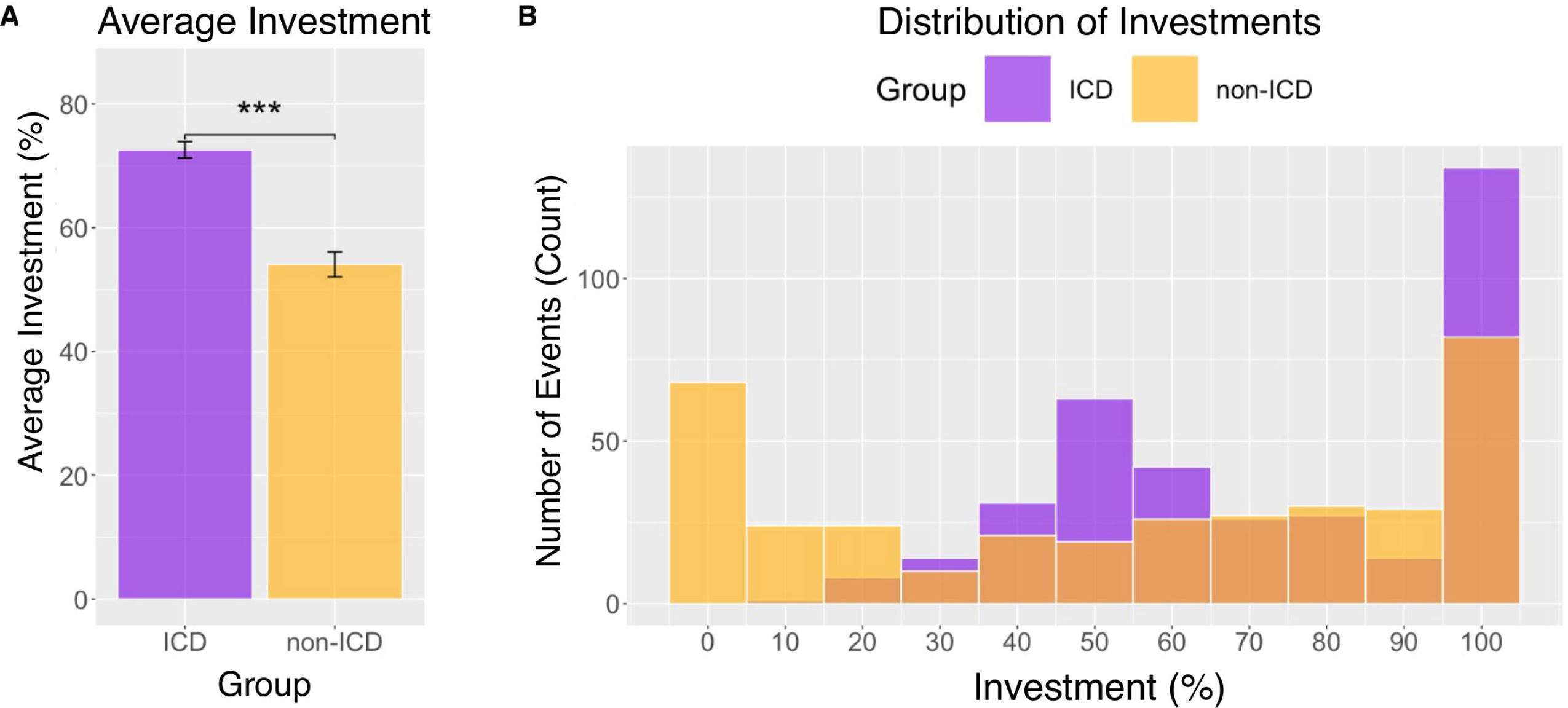
Patients with ICD risk more than non-ICD patients. Investment choices are summarized separately for the ICD group, shown in purple, and the non-ICD group, shown in yellow. (A) The average investments for the ICD and non-ICD groups are shown. The average investment as a percentage (%) of available investment portfolio is on the y-axis. The groups, ICD and non-ICD, are on the x-axis. (B) The distribution of investments for the ICD and non-ICD groups are shown. The number of trials for which participants selected an investment is on the y-axis. The investment as a percentage (%) of available investment portfolio is on the x-axis. ***=Statistically significant group comparison with a p-value less than 0.001.

**Figure 3.**
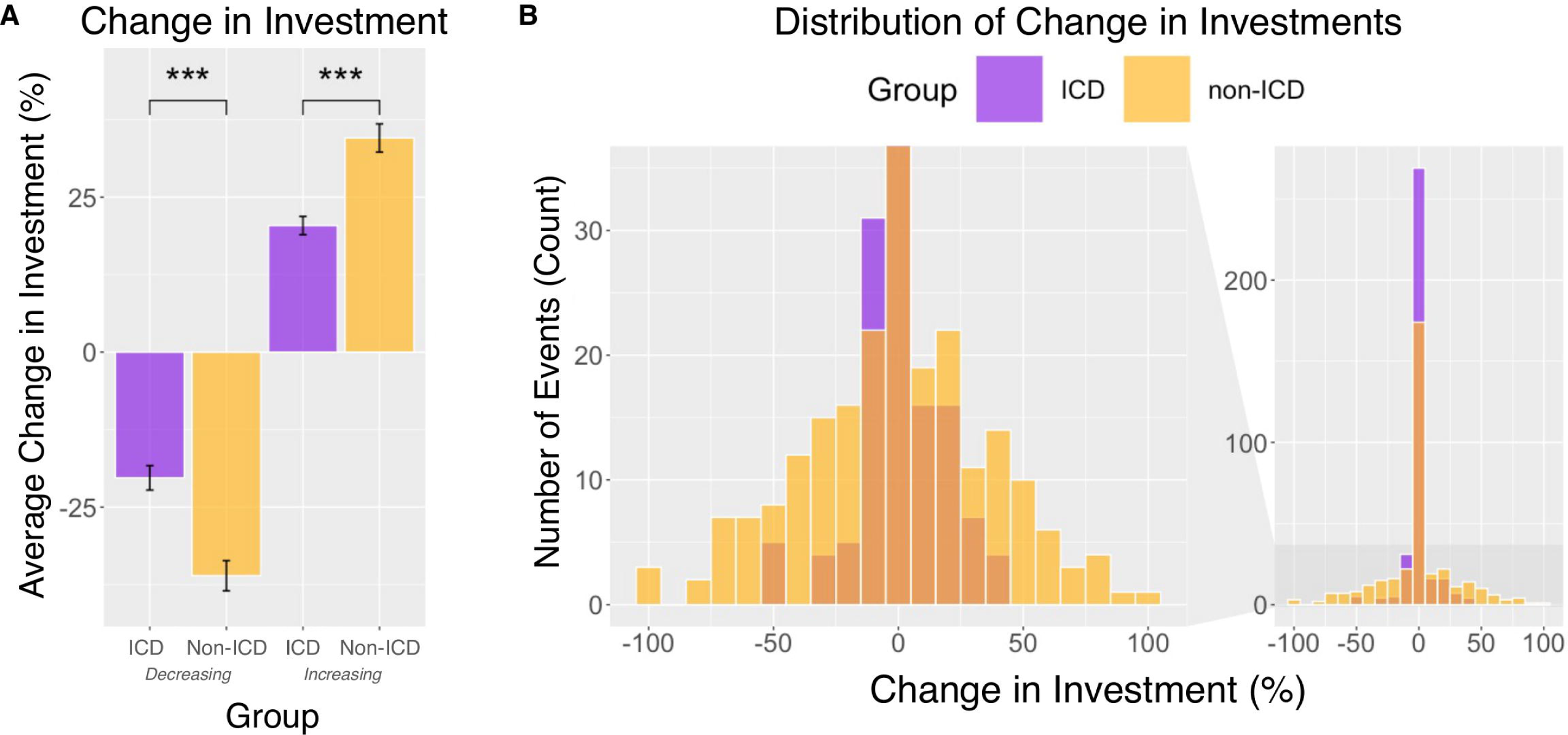
Patients with ICD change their investments less than non-ICD patients. Change in investments are summarized separately for the ICD group, shown in purple, and the non-ICD group, shown in yellow. (A) The average change in investment for the ICD and non-ICD groups are shown. This data is separated by whether participants are decreasing or increasing their investments from the prior trial. The average change in investment as a percentage (%) of available investment portfolio is on the y-axis. The groups are on the x-axis separated by ICD group (ICD versus non-ICD) and investment behavior (decreasing versus increasing investments from the prior trial). (B) The distribution of change in investments for the ICD and non-ICD groups are shown. The number of trials for which participants changed their investment by a specified amount is on the y-axis. The change investment as a percentage (%) of available investment portfolio is on the x-axis.

### Computing Behavioral Variables to Sort Sub-Second Dopamine

For the time period before the outcome reveal (Figure 4), we calculated the change in investment between: the percentage of a participant’s portfolio invested in the current trial, and the percentage of their portfolio invested in the prior trial. For the time period after the outcome reveal (Figure 5), we calculated a reward prediction error (RPE) and a punishment prediction error (PPE) on every trial using the following equations (note: RPEs and PPEs were calculated separately on each trial, following the valence-partitioned reinforcement learning (VPRL) hypothesis^25,26^):

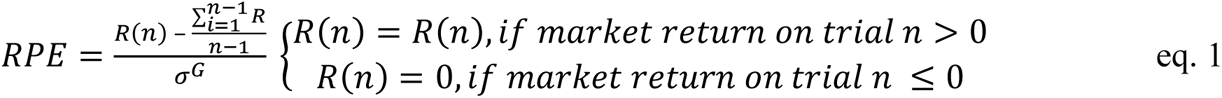

**Figure 4.**
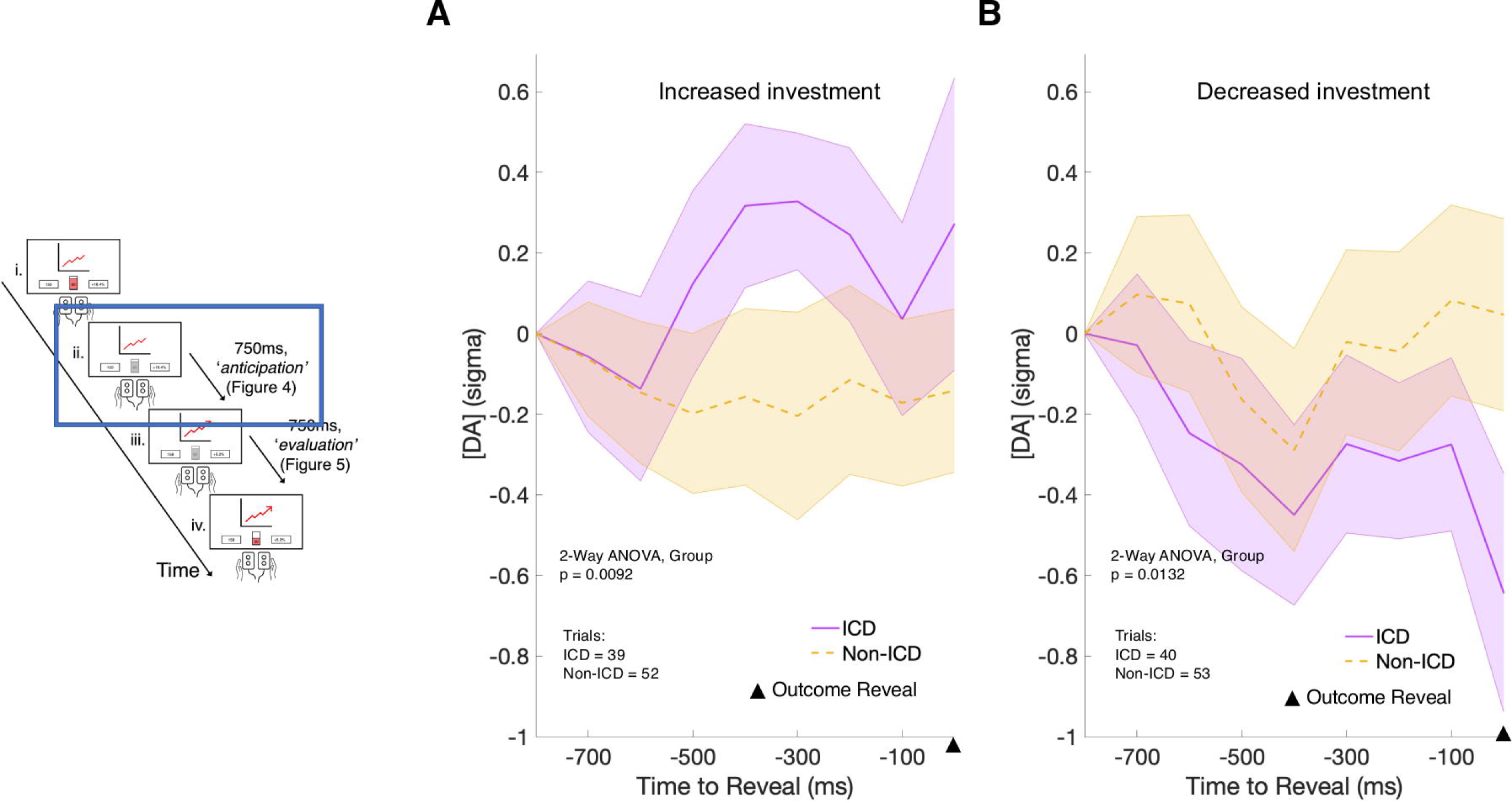
Sub-second dopamine fluctuations in ICD rise with increasing investments and fall with decreasing investments. Sub-second dopamine fluctuations for the time period before the outcome reveal are separated into the ICD group, shown in purple, and the non-ICD group, shown in yellow. The investment outcome reveal is designated on the x-axis with a triangle. A) The average dopamine concentrations on trials for which the investment change is less than 0% to - 30% are shown. The average relative dopamine concentration ([DA]) is on the y-axis. This concentration is normalized to each individual before group-level averaging. The time series is on the x-axis as a countdown until the outcome reveal. B) The average dopamine concentrations on trials for which the investment change is greater than 0% to 30% are shown. The average relative dopamine concentration ([DA]) is on the y-axis. This concentration is normalized to each individual before group-level averaging. The time series is on the x-axis as a countdown until the outcome reveal.

**Figure 5.**
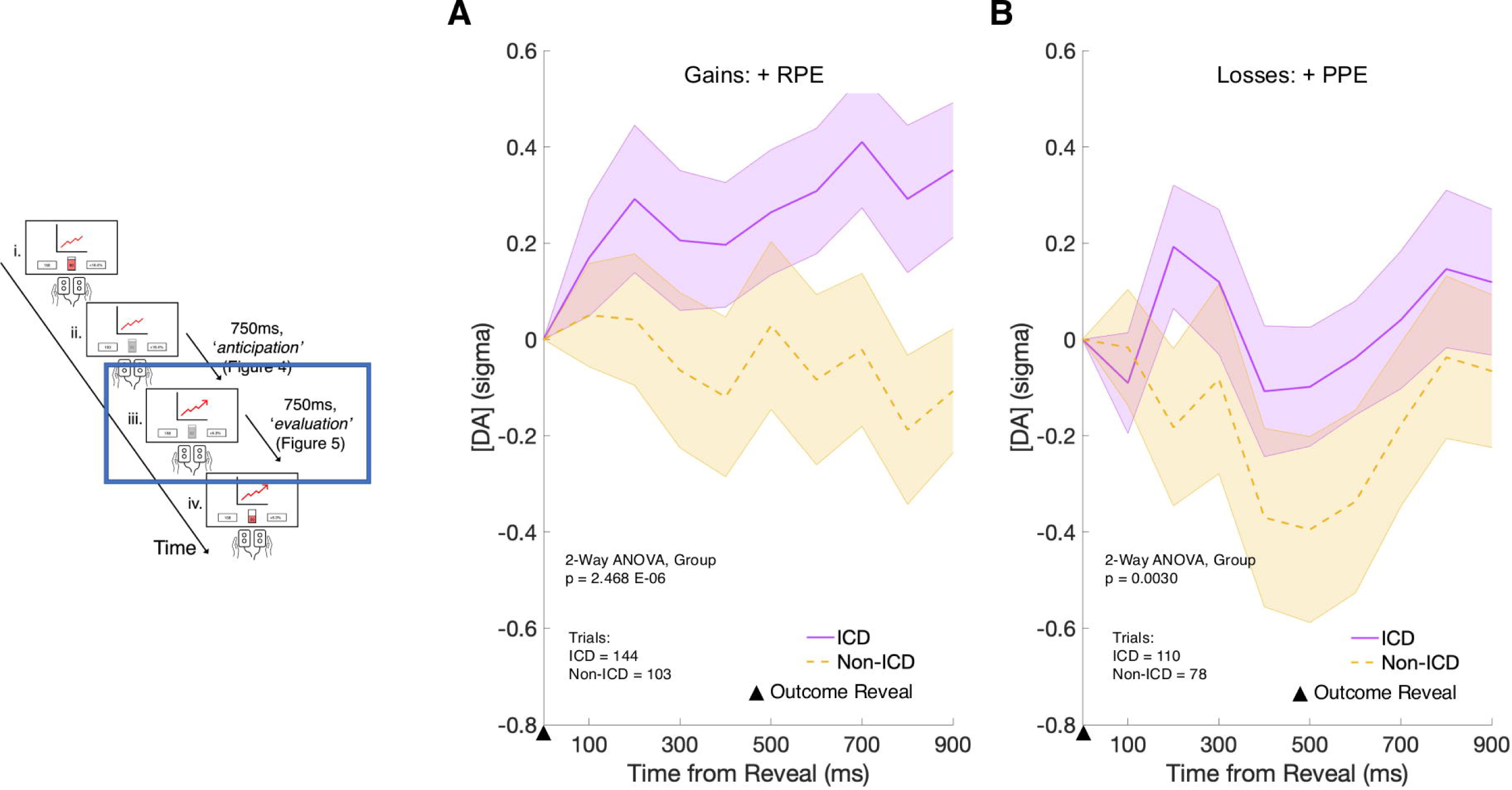
Sub-second dopamine fluctuations increase in ICD after better-than-expected gains and worse-than-expected losses. Sub-second dopamine fluctuations for the time period after the outcome reveal are separated into the ICD group, shown in purple, and the non-ICD group, shown in yellow. The investment outcome reveal is designated on the x-axis with a triangle. A) The average dopamine concentrations on gain trials for which the reward prediction error (RPE) is positive (+) are shown. The average relative dopamine concentration ([DA]) is on the y-axis. This concentration is normalized to each individual before group-level averaging. The time series is on the x-axis starting at the investment outcome reveal. B) The average dopamine concentrations on loss trials for which the punishment prediction error (PPE) is positive (+) are shown. The average relative dopamine concentration ([DA]) is on the y-axis. This concentration is normalized to each individual before group-level averaging. The time series is on the x-axis starting at the investment outcome reveal.

Where *R*(*n*) is the participant’s return on trial “*n*”,*σ*^*G*^ is the standard deviation of participant returns over all previous trials where on each trial returns that were less than zero were treated as null outcomes by the hypothesized reward system^25,26^). Likewise, PPEs were calculated as follows:

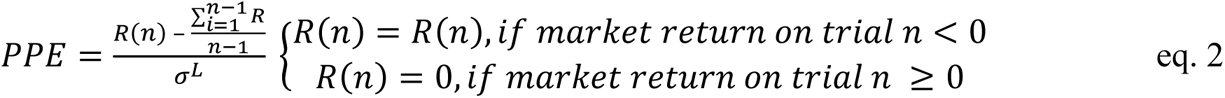

Where *σ*^*L*^is the standard deviation of participant returns over all previous trials where on each trial returns that were greater than zero were treated as null outcomes by the hypothesized punishment system ^25,26^). This partitions the results of each investment decision by the valence of its outcome such that the statistics of each can be assessed independently.^25,26^

The computed behavioral variables are used to sort and compare dopamine fluctuations separately for the time periods before and after the outcome reveal. For the time period before the outcome reveal, the patients’ trial-to-trial investment change was used to sort dopamine fluctuations. This returned three categories for analysis: 1) investment changes less than 0% to - 30%, 2) investment changes equal to 0%, and 3) investment changes greater than 0% to 30%. For the time period after the outcome reveal, dopamine fluctuations were sorted by both gains versus losses and RPE versus PPE.

### Dopamine Measurements

The protocol for dopamine measurements is the same as reported in previous work.^14,16,17,23^ Dopamine measurements were made during the functional mapping phase of DBS electrode implantation surgery. Participants completed the stock market investment task within 30 minutes while electrochemical measurements are made on carbon fiber microelectrodes as previously reported. DBS therapeutic targets and clinically safe trajectories for DBS electrode insertion determine the potential research recording sites. The neurosurgeons on the research team place the carbon-fiber microsensors into the striatum prior to the placement of functional mapping electrodes in the STN or GPi, respectively. The neurosurgeons use one of five potential microelectrode recording trajectories made available by a five-hole “Ben-gun” array attached to the Cosman–Roberts–Wells (CRW) stereotactic headframe.

The carbon-fiber microsensor electrodes are constructed by our laboratory to match the dimensions of the tungsten microelectrodes used for functional mapping during DBS electrode implantation surgery. These custom carbon fiber microelectrodes have passed a successful Ethylene Oxide Sterilization Exposure and Sterility Audit conducted by BioLabs to ensure preoperative ethylene oxide treatment fully sterilizes the carbon-fiber microsensor electrodes. These electrodes have also been validated and approved for autoclave and hydrogen peroxide sterilization. We have compared DBS cases with and without our research protocol and are able to report no increase in infection risk attributable to the research protocol.^18^

We used fast-scan cyclic voltammetry (FSCV) protocol and analysis approach established in prior work.^14,16,17,23^ Carbon-fiber microsensors are conditioned using a 60-Hz application of the triangular measurement waveform for approximately 10 minutes. Immediately following, the measurement protocol is initiated, wherein continuous measurements of current at collected with at 100kHz sampling rate. The measurement protocol consists of a repeating pattern: the electrode potential is held at −0.6 Volts (V) for 90 milliseconds (ms), ramped up to +1.4 V at 400 Volts/second (V/s) for 5 ms, and ramped back down to −0.6 V at −400 V/s for 5 ms. The 1,000 samples of current (maximum range: +/− 2,000nA) collected during the triangular voltage protocol are used to estimate dopamine concentrations. The carbon-fiber microsensor is attached to a portable electrochemistry recording rig.^23^ The recording rig includes a head stage (CV-7B/EC; Axon Instruments), an amplifier (700B; Axon Instruments Multiclamp), an analog-to-digital (A/D) converter (Digidata 1440A; Axon Instruments), and a laptop (MacBookPro; Apple).^23^ A separate laptop (MacBookPro; Apple) is used for task presentation and behavioral data collection. To synchronize the timelines of the electrochemistry and behavioral computers, an independent signal generator (Tektronix AFG320 Arbitrary Function Generator) sends a split signal to both the behavioral computer and the A/D convertor. The signal generator provides an analog square waveform (voltage step 0 to 5V, at 1Hz, with 1% duty cycle) to align the Digidata 1440A data stream with the behavioral data.

### Statistical analyses

We estimated dopamine concentrations from the electrochemical measurements based on the methods and models previously published.^14,16,17,23^ We performed statistical analyses and generated figures using RStudio and MATLAB. We compared dopamine and behavior data across ICD (N=3) and non-ICD (N=3) groups. In our behavior analyses, we compared the ICD and non-ICD groups using a paired-sample t-test for average investment and a two-sample t-test for change in investment. We pooled trial data from all participants in each group prior to group-level characterization of dopamine fluctuations. We analyzed dopamine fluctuations during two time periods: the 800 milliseconds before the outcome reveal (Figure 3) and the 900 milliseconds after the outcome reveal (Figure 4) for each trial. We centered dopamine data in each group to a baseline level relevant to examine the change in dopamine levels in reaction to a task event. We selected a baseline of 800 milliseconds prior to the outcome reveal to assess anticipatory responses in dopamine leading into the revelation of the investment return. We selected a separate baseline at the time of outcome reveal to assess the change in dopamine levels in reaction to the participant winning or losing on their investment. We identified statistically significant differences in dopamine fluctuations using a 2-way analysis of variance (ANOVA) using group (ICD versus non-ICD) and time as independent variables (Supplemental Tables 2-4).

### Data availability

The authors confirm that the data supporting the findings of this study are available within the article and its supplementary material.

## Results

### Participant Characteristics

We collected clinical characteristics for each participant to separate participants into ICD and non-ICD groups (Table 1). All 6 patients were male and receiving DBS for PD and completed our task off their dopaminergic medications. The non-ICD patients were selected to match the ages represented in the ICD group. All ICD patients demonstrated impulsivity following dopamine receptor agonists; likewise, impulsivity symptoms resolved after cessation of these medications.

### Risk behavior in Participants with versus without Impulse Control Disorder

Participants performed the stock market investment task while off all dopaminergic medications during DBS electrode implantation surgery. On average, participants with ICD invested significantly more than the non-ICD group (Figure 2A, paired-sample t-test, p-value=3.8133e-11). Notably, an investment of 100% is the most frequently selected option in the ICD group (Figure 2B). Examination of the distribution of investments for each group revealed that the non-ICD group, compared to ICD group, expressed more variance and distributed their investments more evenly across the available investment options (Figure 2B).

To determine how patients with ICD react to the uncertain stock market returns, we examined trial-to-trial changes in investment levels (Figure 3) in both groups. This analysis reveals that the ICD group makes significantly smaller adjustments in their investments compared to the non-ICD group when increasing (two-sample t-test, p=6.4979e-05, Figure 3A) or decreasing (two-sample t-test, p-value=4.5731e-06, Figure 3A) their investments. Further, the ICD group leaves their investment “as is” near twice as many times as the non-ICD group (Figure 3B). The ICD group demonstrates a much narrower distribution of investment change with a standard deviation of 10.85% versus the non-ICD group standard deviation of 29.58% (Figure 3B). Together, the ICD group invests higher average investments with smaller changes in investments than the non-ICD group.

### Phasic Dopamine Fluctuations in patients with versus without ICD

Participants’ changes in expectations may be expressed by their willingness to increase their investment level when they expect to win and to decrease their investment level when their expectations of winning decrease. Notably, changes in investment levels in the range of −30% to +30% capture 99.7% of the ICD behavioral data, thus we restricted our analysis to this range in both groups for comparison. Following the submission of participants’ investment decision, in each trial, dopamine levels rise (in ‘anticipation’ of the outcome revelation) when ICD participants had increased their investment by a small amount (+10 to +30%), while non-ICD participants dopamine levels maintain level (Figure 4A). The difference in these dopamine responses is significant across ICD group (2-Way ANOVA, p=0.0092). Likewise, dopamine levels fell when the ICD group decreased their investment by a small amount (−10 to −30%), while non-ICD participants dopamine levels dip within 100-200ms before returning to level (Figure 4B). The difference in these dopamine responses is also significant across ICD group (2-Way ANOVA, p=0.0132).

In the second phase of reward processing, market return ‘evaluation’ (Figure 1D-iii), we observe, that dopamine levels rise in the ICD group with positive reward prediction errors when participants win money, but not in participants without ICD (Figure 5A). The difference in the response between ICD and non-ICD groups are significantly different across ICD group (2-Way ANOVA, p=2.468 E-06). When participants lose money and experience punishment prediction errors, dopamine levels fall in the non-ICD group (Figure 5B). However, the dopamine response in the ICD group shows a rapid phasic increase before returning and maintaining level (Figure 5B). The difference in the response between ICD and non-ICD groups are significantly different across ICD group (2-Way ANOVA, p=0.0030).

## Discussion

We report the first direct measurements of dopamine levels in the human brain with sub-second temporal resolution from patients with a history of Impulse Control Disorder caused by dopamine receptor agonist therapy.^14,16,17,23^ We compare these measurements to similar measurements from patients without a history of ICD. These data were obtained while all patients were off of all prescribed dopaminergic medications, which provides a launching point for future pre-clinical investigations of dopamine’s role in ICD. Volunteers undergoing a DBS electrode implantation procedure for PD treatment participated in an IRB approved research procedure with patient consent obtained at the pre-operative visit.^18^ Participants performed a sequential stock market investment task^16,23,24^ allowing the investigation of risk-taking behavior, changes in behavior due to consequences of recent actions, and the action of moment-to-moment changes in dopamine levels in anticipation and in reaction to expected and unexpected outcomes.

In our cohort, patients with ICD invest significantly more than age- and sex-matched patients undergoing the same procedure (Figure 2). Furthermore, patients with ICD do not change their exposure to risk as much as controls (Figure 3). Dopamine levels rapidly rose in anticipation of outcomes in patients with ICD, but not controls (Figure 4A). In anticipation of losses, dopamine levels fell more dramatically in patients with ICD (Figure 4B). Then, following ‘better than’ or ‘worse than expected’ outcomes, dopamine levels rapidly rose or fell, respectively; however, patients with ICD demonstrate a response that is biased toward positive reinforcement of risky actions (Figure 5). Together, our results suggest that patients with PD and a history of ICD possess a dopaminergic system that may be amplified in its response to risk related stimuli and outcomes; and, thereby more prone to find these stimuli, actions, and associated outcomes positively reinforcing. This pattern is particularly profound given its identifiable presence on gameplay even in the absence of the dopamine receptor agonists that are usually required to clinically recognize ICD and its harmful symptoms. We hypothesize that future pre-clinical studies may clarify reproducible behavioral signatures that correlate to an underlying propensity to engage in risky actions.

Dopamine fluctuations in anticipation of outcomes track behavior that is consistent with changes in expectations in our stock market task. A rational player in the stock market task will increase their investment when their expectations of a positive outcome are also increased; similarly, they will decrease their investment when their expectations of a positive outcome decrease. The logic in this task was designed to follow common sense: if a player thinks they have a higher chance of winning, they will bet a higher amount; similarly, if a player thinks they have a higher chance of losing, they will bet a lower amount. Dopamine fluctuations within 800 milliseconds of an anticipated outcome clearly differentiate changes in patients’ investment levels. Patients without a history of ICD show a relatively flat response following small increases or decreases in their investment consistent with a muted reaction to expressed expectations. Whereas patients with ICD show an amplified reaction in this phase of the trial. In both patient groups, a decrease in investment levels, consistent with an overall decrease in expectations of a positive outcome, resulted in a drop in dopamine levels. However, this signal was exaggerated in the ICD group compared to the non-ICD group. Such a drop in dopamine levels can be interpreted as a negative reinforcer of behavior, which is consistent with ICD patients being less likely to decrease their investment exposure.

The present study brings together work using sub-second dopamine fluctuations identified on a behavioral task to separate addiction pathology with evidence of this dopamine originating in the dorsal striatum, namely the caudate and putamen. Our group recently demonstrated that sub-second dopamine fluctuations in response to calculated relief values effectively separated patients with and without Alcohol Use Disorder (AUD).^19^ Here we expand this approach to propose sub-second dopamine fluctuations from the dorsal striatum as potential future investigative anatomic structures for preclinical work on the origins of ICD. Our sample size was modest (n = 3 patients with ICD, n = patients without ICD), but our results align with previous human neuroimaging studies demonstrating dorsal striatal activation during risky decision-making in patients with ICD.^10–12^ The insights gained by presenting sub-second dopamine fluctuations measured in the human operating room offer potential mechanisms underlying this debilitating behavioral addiction disorder for future neuroimaging and other pre-clinical work.^19^

This study reports the first measurements of dopamine release with sub-second temporal resolution in patients with a history ICD. A major challenge for future work utilizing approaches like those described here will be to determine to what extent findings in these relatively restricted patient populations may or may not generalize to other patient populations or to more generalizable brain mechanisms supporting human thoughts, feelings, and actions. An additional challenge will be recruiting a larger sample size to confirm our findings, given our measurement method does not allow patient selection by pathology. However, the profile of dopaminergic activity identified within the ICD group may have more immediate translational potential (following additional work) as potential functional targets for closed-loop DBS therapeutic intervention for use in addiction disorders.^27,28^ Our results demonstrate that intracranial measurements from DBS to treat movement disorders can provide insight into neuropsychiatric pathologies like impulse control disorder and related addiction disorders.

## Supplementary material

Supplementary material is available at *Brain* online.

## Supporting information

Dopamine in Impulse Control Disorder Supplement

