## Supplementary material for "Sub-second Dopamine Signals during Risky Decision-Making in Patients with Impulse Control Disorder": Dopamine in Impulse Control Disorder Supplement

475 Vine St.

Winston-Salem, NC 27101

**Running title**: Dopamine in Impulse Control Disorder

### Supplemental Figures:

#### Table 1:

***Research Electrode Planning Coordinates***

| *Patient* | Location | Midcommissural Point Coordinates (mm) | | |
|---|---|---|---|---|
| | | Lateral | Anterior-Posterior | Superior-Inferior |
| *1* | Putamen | 21.4694 | 10.8178 | 10.1088 |
| *2* | Caudate | 15.2602 | 4.0493 | 20.7628 |
| *3* | Caudate | 13.8364 | 12.2779 | 22.1552 |
| *4* | Caudate | 17.3988 | 11.0097 | 17.2557 |
| *5* | Caudate | 16.249 | 11.8752 | 18.7715 |
| *6* | Caudate | -16.0452 | 13.4861 | 21.5069 |

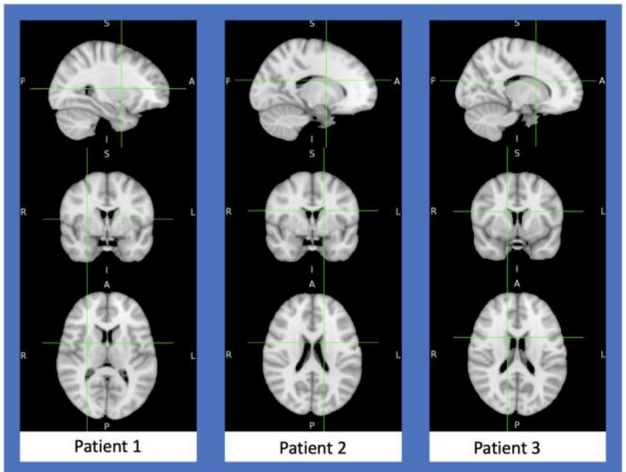


Supplementary Figure 1a: Columns 1-3, left to right correspond to patient's 1, 2, & 3 voltammetry recording electrode's site normalized to the MNI 152 1 mm template. Rows 1-3, top to bottom, display the sagittal, coronal, and axial plane for each patient, respectively. Patients 1-3 all belong to ICD group.

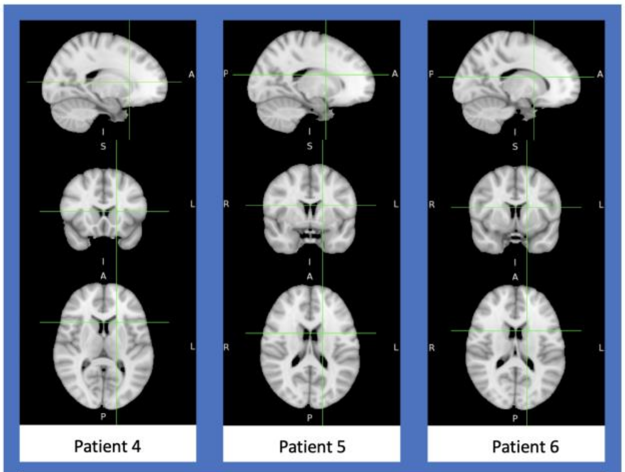


Supplementary Figure 1b: Columns 1-3, left to right correspond to patient's 4, 5, & 6 voltammetry recording electrode's site normalized to the MNI 152 1 mm template. Rows 1-3, top to bottom, display the sagittal, coronal, and axial plane for each patient, respectively. Patients 4-6 all belong to non-ICD group.

Note: We used FSL[1] to map electrode coordinates to a standard Montreal Neurological Institute (MNI) template.[2]

1. Jenkinson M, Beckmann CF, Behrens TEJ, Woolrich MW, Smith SM. FSL - Review. *Neuroimage*. Published online 2012.
2. Collins DL, Zijdenbos AP, Kollokian V, et al. Design and construction of a realistic digital brain phantom. *IEEE Trans Med Imaging*. Published online 1998. doi:10.1109/42.712135

#### Table 2:

**2-Way ANOVA Comparing ICD versus Non-ICD Groups: Leading into the Reveal**

| | Context | | |
|---|---|---|---|
| Variable | DA: < 0% to -30% Investment Change | DA: 0% Investment Change | DA: > 0% to 30% Investment Change |
| Time | 0.756922793 | 0.956370803 | 0.96231549 |
| Group | 0.01321199 | 0.088254828 | 0.009193345 |
| Time*Group | 0.95694099 | 0.671807815 | 0.900070348 |

#### Table 3:

**2-Way ANOVA Comparing ICD versus Non-ICD Groups: Gains, Post-Reveal**

| | Context | |
|---|---|---|
| Variable | DA: + Reward Prediction Error | DA: - Reward Prediction Error |
| Time | 0.980948491 | 0.804084585 |
| Group | 2.48303E-06 | 0.10901517 |
| Time*Group | 0.952446668 | 0.986834235 |

### Table 4:

**2-Way ANOVA Comparing ICD versus Non-ICD Groups: Losses, Post-Reveal**

| | Context | |
|---|---|---|
| Variable | DA: - Punishment Prediction Error | DA: + Punishment Prediction Error |
| Time | 0.923836734 | 0.309636526 |
| Group | 0.304677732 | 0.002970958 |
| Time*Group | 0.915156743 | 0.95313243 |
